# Eliciting a potent antitumor immune response by expressing tumor antigens in a skin commensal

**DOI:** 10.1101/2021.02.17.431662

**Authors:** Y. Erin Chen, Katayoon Atabakhsh, Alex Dimas, Kazuki Nagashima, Michael A. Fischbach

## Abstract

Immune modulation has become central to treating cancer. However, global immune stimulation is only effective in a subset of patients and can lead to serious complications, including colitis and type I diabetes. Newer modalities like engineered T cells and tumor vaccines are more specific, but they have shown limited efficacy in solid tumors and are difficult to scale. Bacterial strains from the human microbiome can induce antigen-specific T cells to help maintain barrier function. Here, we redirect CD8+ and CD4+ T cells elicited by the skin commensal *Staphylococcus epidermidis* to recognize tumor cells by expressing tumor-derived antigens in the bacterial cell. *S. epidermidis* expressing the model antigen ovalbumin (*S. epidermidis-*OVA) stimulates antigen-specific CD8+ and CD4+ T cells in vitro. The subcellular localization of the antigen skews the response: cell wall-attached OVA preferentially stimulates CD8+ T cells whereas secreted OVA predominantly induces CD4+ T cells. In a syngeneic tumor model (OVA-expressing B16 melanoma), mice colonized topically with *S. epidermidis-*OVA exhibit a marked reduction in subcutaneous tumor volume compared to mice colonized with *S. epidermidis* expressing mCherry; this effect is dependent on live bacteria and a combination of CD8+ and CD4+ T cells. *S. epidermidis-*OVA also reduces tumor burden when tumor cells are injected intravenously (a model of metastasis), demonstrating that the antitumor effect operates in tissues distant from the site of bacterial colonization. *S. epidermidis* strains expressing neoantigen peptides from the B16 tumor cell line exhibit potent antitumor efficacy without inducing an autoimmune response against melanocytes in healthy tissue. Antigen-expressing colonists are a simple but powerful strategy to elicit a targeted T cell response in the context of cancer and other diseases.

## MAIN TEXT

Immune modulation has become a central component of cancer therapies, with inhibitors of the checkpoint proteins PD-1 and CTLA-4 most widely deployed^1^. Although checkpoint blockade is efficacious across many cancer types, it only works in a subset of patients. Moreover, global stimulation of immune function frequently induces autoimmunity (e.g., colitis and type 1 diabetes)^2,3^. Thus, a central challenge in immuno-oncology is to develop methods for stimulating immune cells that recognize cancer cells selectively.

There are two predominant strategies for eliciting specific immune responses in the context of oncology. The first is to generate antigen-specific T cells by ex vivo transduction or electroporation with a chimeric antigen receptor (CAR) or a T cell receptor (TCR)^4,5^. This approach has shown great promise in treating hematologic malignancies but has several downsides: it is expensive^6^, engineered T cells can be prone to exhaustion^7^, and there have been challenges in getting CAR-T cells to act against solid tumors^8^. The second strategy is to vaccinate the host with tumor neoantigens or antigen-loaded dendritic cells; these approaches have yielded promising results in preclinical models but are not yet consistently efficacious in clinical trials^9–12^.

Certain strains of the gut and skin microbiota induce antigen-specific T cells^13–16^, raising the possibility of a simpler way to elicit a targeted T cell response that does not require ex vivo cell engineering. However, the T cells induced by the microbiota harbor TCRs specific for bacterial antigens, limiting their utility in treating cancer or autoimmune disease. Here, we show that this process can be redirected to elicit antigen-specific T cells against non-bacterial antigens. By expressing a model antigen or a tumor-derived neoantigen in the T cell-stimulatory skin commensal *Staphylococcus epidermidis* LM087, we elicit tumor-specific CD8+ and CD4+ T cells. These cells protect against local and metastatic progression of a poorly immunogenic mouse melanoma.

### Selecting *Staphylococcus epidermidis* LM087 as the tumor antigen chassis

Previous efforts to engineer a bacterial strain to elicit an antitumor response have involved the pathogens *Listeria monocytogenes* and *Salmonella typhimurium*. In both cases, antigen-specific T cell responses leading to tumor regression have been observed^17,18^. However, the antitumor efficacy of *Listeria-* and *Salmonella-*based vaccines relies on nonspecific immune activation resulting from local tissue damage, infection and replication within antigen-presenting cells, or direct infection of tumor cells^19^. Efforts have been made to attenuate each pathogen to alleviate the toxicity of infection, but attenuation typically dampens the desired immune response^19^. Efforts to use non-pathogenic *Escherichia coli* strains to boost antitumor immunity also hold promise, but require intratumoral delivery^20,21^.

We decided instead to work with a prevalent member of the healthy human skin microbiome. *Staphylococcus epidermidis* LM087 induces antigen-specific CD8+ T cells in mice and non-human primates^15,22,23^. Importantly, it does so in the context of physiologic skin colonization—without mounting an infection, breaching the skin barrier, or causing any pathologic response. This strain colonizes the skin for an extended period of time after a single application; the CD8+ T cell response it elicits in mice is durable for nine months^22^. We hypothesized that *S. epidermidis* could be a promising starting point for eliciting antitumor immunity given its lack of toxicity and the specificity and potency of the CD8+ T cell response observed in mice and non-human primates following topical application.

### Developing a genetic system for *S. epidermidis*

Despite the fact that *S. epidermidis* colonizes the skin of every human, methods to manipulate it genetically are poorly developed; only a handful of mutants have ever been reported, and only in strains that adsorb phage efficiently or are domesticated^24,25^.

Targeted genetic modification of *S. epidermidis* has been challenging for two reasons: *S. epidermidis* has multiple stringent restriction systems that differ substantially among strains^24,26,27^, and— as with many other Gram-positive bacteria—electroporation is an inefficient means of introducing DNA, owing to the thick cell wall^28^. To bypass poor electroporation efficiency, a genetic approach was recently described that involves transduction with bacteriophage Φ187 from an engineered restriction-deficient S. *aureus* (PS187 Δ*hsdR* Δ*sauPSI*) to *S. epidermidis* or other coagulase-negative *Staphylococcus* species on the basis of their similarity in wall teichoic acid structure^29,30^. Although this method works well for a specific clade of *S. epidermidis* strains, it is time-consuming, requires prolonged phage propagation, and is useful only for certain strains of *S. epidermidis* with efficient phage adsorption capacity. Given the high variability of phage specificity across *S. epidermidis* strains, there are many primary isolates of *S. epidermidis* for which this method does not work.

Reasoning that antigen engineering would require an efficient method for editing the genomes of primary isolates, we started by developing a new genetic system for *S. epidermidis*. By integrating elements of electroporation-based protocols for Gram-positive bacteria and yeast^31–34^, we developed a protocol with three key elements: to prepare electrocompetent cells, we cultivate *S. epidermidis* in hyperosmolar sorbitol, washing thoroughly in high volume 10% glycerol; to prepare the DNA, we pass the plasmid through a the methylase-deficient (Δ*dcm*) *E. coli* strain DC10B^24^; and directly before (Method 1) or after (Method 2) electroporation, we introduce a heat shock (**Figure 1A**). Using this method, we were able to construct mutations in 13 of the 17 *S. epidermidis* strains we tested, include >10 primary human isolates from diverse phylogenetic groups (**Figure 1B**). Our approach does not restrict the efficiency of cloning to phage type and does not require specialized knowledge of strain-specific restriction systems.

**Fig. 1.**
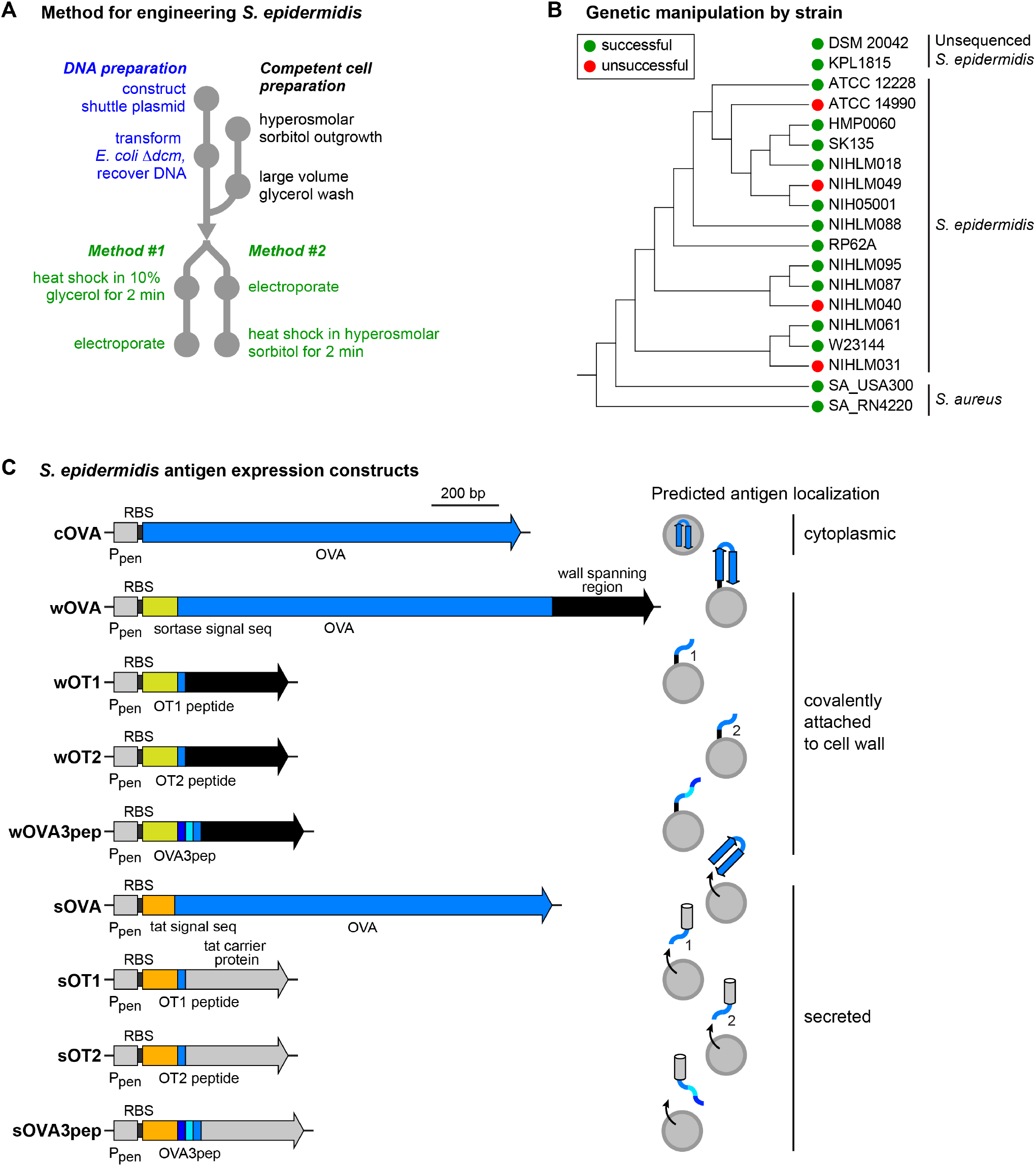
Engineering *S. epidermidis* to express non-native antigens. (A) Schematic of the new genetic system for *S. epidermidis*. To bypass stringent restriction systems, the target plasmid is passed through *E. coli* Δ*dcm*. To optimize cell competency, *S. epidermidis* is grown in media containing hyperosmolar sorbitol, harvested during late-log phase, and thoroughly washed with a large volume of 10% glycerol to eliminate salts. *Staphylococcus* is subjected to heat shock before (Method #1) or after (Method #2) electroporation with the plasmid. (B) Phylogenetic tree of *S. epidermidis* and *S. aureus* strains. This method was successful in 13 of the 17 primary *S. epidermidis* isolates in which we tested it, including two strains that do not have genome sequences available in NCBI. Green = genetically manipulatable. Red = not genetically manipulatable with our method. (C) We constructed strains in which full-length ovalbumin or an ovalbumin-derived peptide is expressed in the cytoplasm, fused to the cell wall, or secreted. The predicted subcellular localization of each antigen is shown at right. Expression is driven by the constitutive *S. aureus* promoter Ppen (gray box) and the ribosome binding site (RBS) from *S. aureus hld*. Ovalbumin (blue box) is expressed either as a full-length protein (OVA), the MHC I-restricted antigenic peptide (OT1), the MHC II-restricted antigenic peptide (OT2), or a concatemer of computationally predicted H2-M3-restricted antigenic peptides (OVA3pep). For localization to the cell wall, the antigen was inserted between the signal sequence from *S. aureus* protein A (yellow box) and a wall-spanning domain that is covalently anchored to the cell wall by sortase (black arrow). For secretion, full-length OVA without its N-terminal methionine is fused to the Tat signal sequence of *S. aureus fepB* (orange box) or an antigen-containing peptide is inserted into *fepB* immediately after the N-terminal signal sequence.

### Engineering antigen expression into *S. epidermidis*

We used this system to engineer *S. epidermidis* LM087 (hereafter, *S. epidermidis*) to express non-native antigens, with the goal of redirecting T cell responses against an antigen of choice. Notably, the process by which *S. epidermidis* antigens are presented to CD8+ T cells is not well understood: the identity of the antigen-presenting cell is unclear, the mechanism by which antigen is presented on class I MHC (active transport from the cytosol vs. cross-presentation) is unknown, and the process that determines which antigens are selected from among thousands of proteins in the bacterial cell has not been studied. Additionally, it was unclear whether a non-native protein could compete against native *S. epidermidis* antigens for CD8+ T cell recognition, and if the foreign antigen would require presentation by the nonclassical MHC Ib molecule H2-M3, which presents immunodominant native *S. epidermidis* antigens to CD8+ T cells^15^.

Our strategy took these elements of uncertainty into account (**Figure 1C**). We started with ovalbumin (OVA) as a model tumor antigen since it harbors well-characterized antigenic peptides that are recognized by OVA-specific CD8+ or CD4+ T cells (OT-1 or OT-2, respectively). In light of the inherent challenges in expressing a non-native antigen in an undomesticated human isolate at high enough levels to be efficiently presented on MHC upon engulfment, we used a *Staphylococcus* replicative plasmid with a constitutive promoter (pLI50-Ppen)^35^ and added the ribosome binding site from the *S. aureus* delta-hemolysin (*hld*) gene, which promotes strong, constitutive translation in *S. aureus* and *S. epidermidis*^36,37^. We designed four forms of the OVA antigen: the full-length protein (OVA), an MHC I-restricted antigen from OVA (amino acids 257-264, here termed ‘OT1’), an MHC II-restricted antigen from OVA (amino acids 329-337, here termed ‘OT2’), or a concatemer of three computationally predicted H2-M3-binding peptides from OVA (‘OVA3pep’)^38^. All forms of the OVA antigen were codon-optimized for *Staphylococcus*.

Next, we generated three sets of strains in which we varied the nature of the antigen and its subcellular localization within *S. epidermidis*: (*i*) One strain for cytoplasmic expression consisting solely of full-length OVA (cOVA). (*ii*) Four strains for cell-wall-displayed antigen in which OVA, OT1, OT2, or OVA3pep is spliced between the *N*-terminal sortase signal peptide and *C*-terminal cell wall-spanning regions of *S. aureus* protein A, yielding *S. epi*-wOVA, *S. epi*-wOT1, *S. epi*-wOT2, and *S. epi*-wOVA3pep. A similar approach has been used for surface display of recombinant proteins in *Staphyloccocus xylosus*^39^. (*iii*) Eight strains for antigen secretion. In six of these strains, OT1, OT2, or OVA3pep are spliced into the secreted proteins FepB (Tat pathway)^40,41^ or SERP0318 (Sec pathway) ^42^ at the predicted signal sequence cleavage site; in the remaining two, full-length OVA is fused to the *N*-terminal Tat or Sec signal sequences from the same proteins. The OVA-FepB chimeras yielded strains *S. epi*-sOT1, *S. epi*-sOT2, *S. epi*-sOVA3pep, and *S. epi*-sOVA; the OVA-SERP0318 chimeras expressed poorly and were not used further.

We verified production of the full-length OVA constructs by Western blot (**Figure S1**). Notably, although *S. epidermidis* has a well-described Sec secretion system and no discernible Tat secretion system, the Tat signal peptide enabled efficient production and secretion of OVA.

### Engineered strains of *S. epidermidis* stimulate antigen-specific T cells in vitro

To test whether the engineered strains of *S. epidermidis* are capable of generating an antigen-specific CD8+ or CD4+ T cell response, we used an in vitro mixed lymphocyte assay in which the bacterial strain is mixed with murine dendritic cells and transgenic OT-1 or OT-2 T cells that are specific for an OVA peptide-MHC complex (**Figure 2A, S2**). We measured T cell activation after four hours of co-culture by monitoring the level of Nur77, an early and specific marker of T cell receptor signaling^43^.

**Fig. 2.**
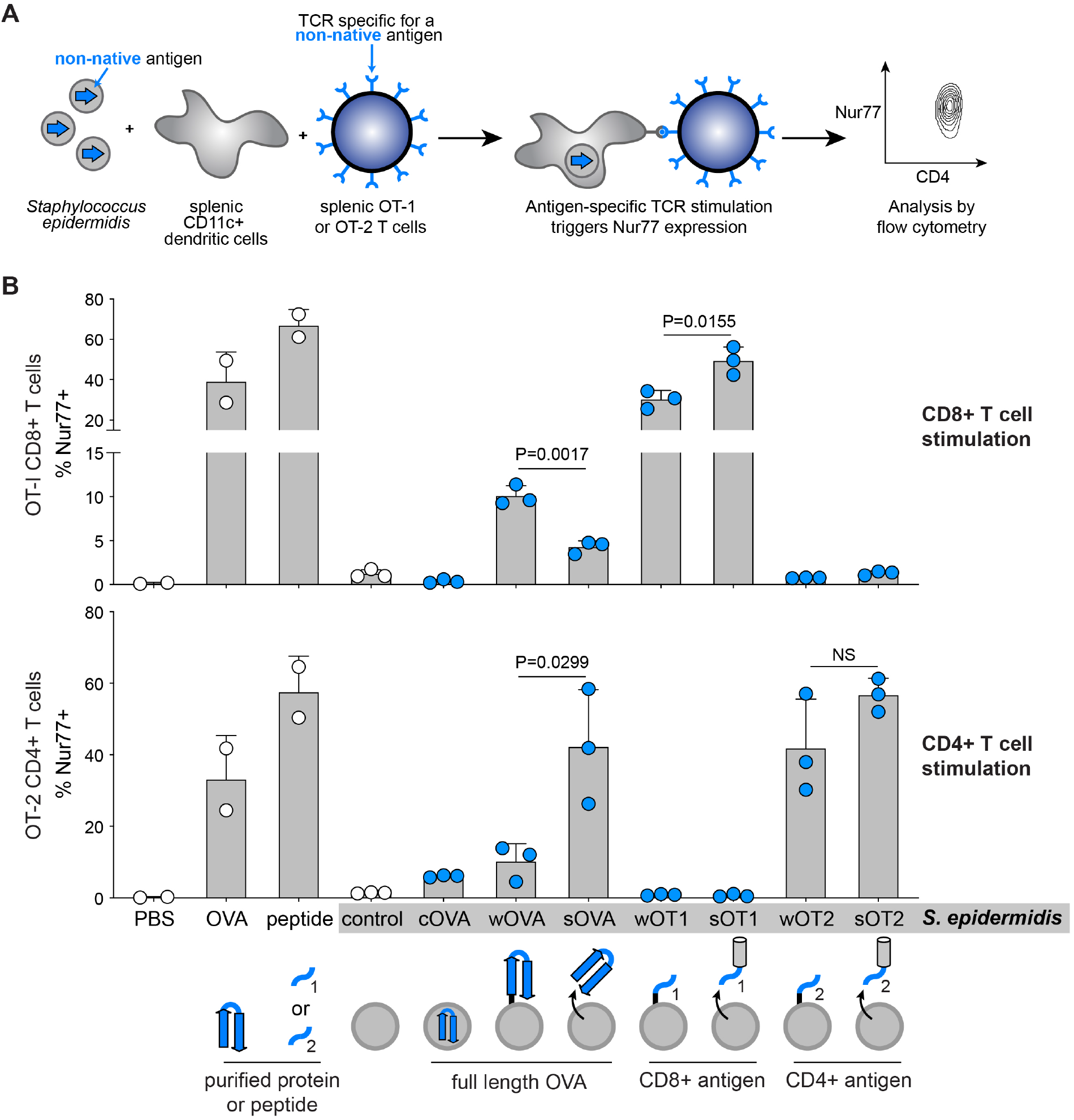
Engineered strains of *S. epidermidis* stimulate antigen-specific T cells in vitro. (A) Schematic of the in vitro assay of T cell activation. Each *S. epidermidis* strain is heat-shocked to prevent bacterial overgrowth of mammalian cells and co-cultured with splenic CD45.1+ CD11c+ dendritic cells for two hours to allow for antigen processing. Transgenic T cells specific for an OVA-derived CD8+ (OT-1) or CD4+ (OT-2) T cell epitope are then added and co-cultured for four hours to enable antigen presentation from dendritic cells to T cells and TCR signaling, which upregulates Nur77 expression. Cells are then placed on ice, fixed, stained for surface markers and intracellular markers, and then analyzed by flow cytometry. T cells are gated on live CD45.1-CD90.2+ TCRβ+ CD8β+ or CD4+ and analyzed for expression of Nur77, an early marker of TCR signaling. (B) Percentage of Nur77+ CD8+ T cells (top) or Nur77+ CD4+ T cells (bottom) in the presence of PBS, purified ovalbumin (OVA), purified OT1 (top) or OT2 (bottom) peptide, or engineered *S. epidermidis* strains (highlighted in gray bar). Representative flow plots are shown in **Figure S1**.

From these data, we draw two conclusions: First, several of our engineered strains stimulate antigen-specific CD8+ and CD4+ T cells robustly, demonstrating that a non-native protein can be expressed at a high enough level in this undomesticated commensal to be presented by MHC I and II (**Figure 2B**).

Second, the subcellular localization of the antigen can dictate a preference toward CD8+ versus CD4+ T cell stimulation. The strain expressing cell wall-attached OVA (*S. epi*-wOVA) induces OVA-specific CD8+ T cells (OT-1) more efficiently than secreted OVA (*S. epi*-sOVA); in contrast, *S. epi*-sOVA preferentially stimulates OVA-specific CD4+ T cells (OT-2) (**Figure 2B**). The strain expressing cytoplasmic OVA (*S. epi*-cOVA) activates CD8+ and CD4+ T cells weakly even though cOVA is expressed at comparable levels to sOVA (**Figure S1**). Notably, strains expressing an OVA peptide rather than the full-length protein (*S. epi*-wOT1, *S. epi*-wOT2, *S. epi*-sOT1, and *S. epi*-sOT2) stimulate CD8+ or CD4+ T cells at high levels regardless of antigen localization.

### *S. epidermidis*-OVA slows the progression of subcutaneous melanoma

Next, we tested the ability of the engineered *S. epidermidis* strains to limit the growth of tumor cells in a syngeneic model of murine melanoma (**Figure 3A**). Given our uncertainty about which T cell subtypes would be involved in antitumor immunity, wild-type specific pathogen free (SPF) C57BL/6 mice were colonized with a combination of *S. epi-*wOT1 and *S. epi-*sOVA (hereafter, ‘*S. epi*-OVA’). Bacterial strains were transferred by gentle topical application to the top of the head using a cotton swab; as established previously, this procedure does not breach the skin barrier but leads to robust skin colonization by live bacteria ^23^. Mice were colonized starting seven days before we subcutaneously injected cells from a C57BL/6-derived melanoma line that expresses ovalbumin, B16-F0-OVA, into the right flank. We did not administer additional immune adjuvant therapy (e.g., checkpoint blockade, cytokines, or adoptively transferred T cells). An *S. epidermidis* strain producing a control protein, mCherry, from the same plasmid backbone was applied topically as a control (*S. epi-*control).

**Fig. 3.**
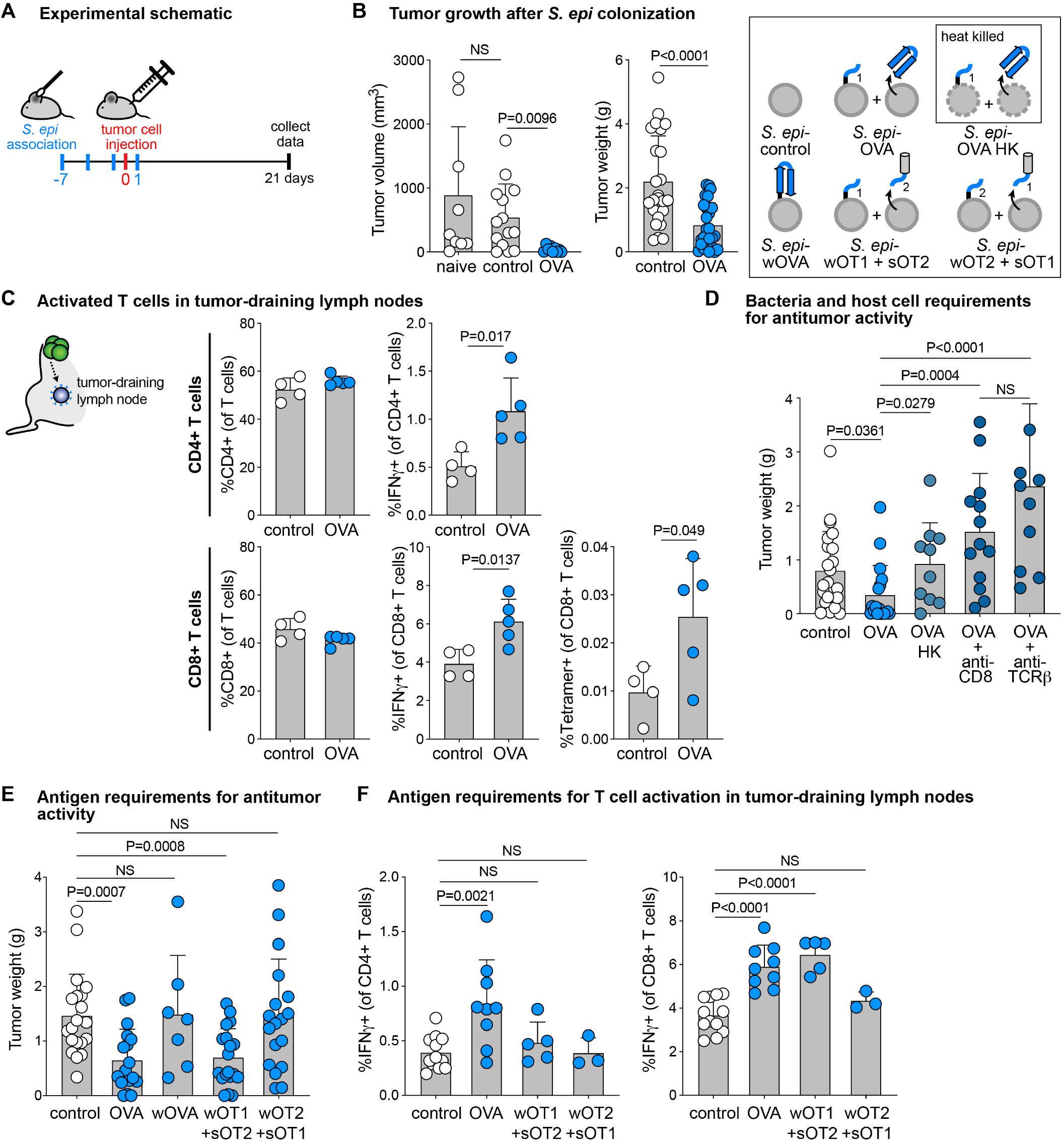
Engineered *S. epidermidis* strains slow tumor progression and stimulate antigen-specific T cells in vivo. (A) Schematic of subcutaneous tumor experiments. Mice are colonized topically by gentle application of live *S. epidermidis* strains to the head with a cotton swab starting seven days prior to tumor injection. On day 0, B16-F0-OVA melanoma cells are freshly prepared from growing cultures and injected subcutaneously into the right flank. Mice are sacrificed on day 21, and tumors and tumor-draining inguinal lymph nodes are collected for analysis. (B) Left panel: day 19 caliper measurements of subcutaneous B16-F0-OVA tumors in naïve SPF mice and mice colonized with *S. epi*-control or *S. epi*-OVA (a 1:1 mixture of *S*.*epi-*wOT1 and *S. epi*-sOVA). Right panel: day 21 masses of dissected subcutaneous tumors. (C) Flow cytometry analysis of tumor-draining inguinal lymph nodes. Frequencies of CD4+ (top) and CD8+ (bottom) cells within live CD90.2+ TCRβ+ T cells do not differ between control (white dots) and *S. epi-*OVA-associated (blue dots) mice, but frequencies of IFN-γ+ cells increased within both CD4+ T cells (top) and CD8+ T cells (bottom). The frequency of MHC I-SIINFEKL tetramer+ cells within CD8+ T cells (bottom right) is increased in *S. epi*-OVA-colonized mice compared to control. (D) Masses of subcutaneous B16-F0-OVA tumors on day 21, dissected from *S. epi-*associated mice. *S. epi*-OVA-colonized mice were treated 2x/week with intraperitoneal injections of 200 μg/mouse anti-CD8α (2.43) or anti-TCRβ (H57-597) neutralizing antibodies (dark blue dots). (E) Masses of subcutaneous B16-F0-OVA tumors on day 21 from *S. epi-*colonized mice. (F) Flow cytometry analysis of tumor-draining inguinal lymph nodes from mice shown in panel E. Frequencies of IFNγ+ cells within live CD90.2+ TCRβ+ CD4+ (left) or CD8+ (right) T cells are shown. Data shown are representative of multiple independent experiments.

*S. epi*-OVA elicited a marked reduction in tumor growth (**Figure 3B**). Nonspecific CD8+ T cell induction by the control strain had no apparent effect; tumors grew just as quickly in *S. epi-*control-treated mice compared to naïve SPF mice (**Figure 3B**). Additionally, although the overall levels of CD4+ and CD8+ T cells are comparable in the tumor-draining lymph nodes, the percentage of IFNγ-expressing CD4+ and CD8+ T cells increase following colonization with *S. epi-*OVA but not *S. epi*-control (**Figure 3C**). The tumor-draining lymph nodes also contain an increased percentage of OVA-specific CD8+ T cells as measured by H2-Kb/SIINFEKL tetramer staining (**Figure 3C**). These results suggest that *S. epi-*OVA elicits an antitumor immune response under conditions of physiologic colonization. Moreover, OVA-expressing colonists induce activated, antigen-specific CD4+ and CD8+ T cells that migrate to the tumor.

Three additional results begin to clarify cellular requirements for the host and microbial colonist. First, heat-killed *S. epi*-OVA failed to stimulate an antitumor response (**Figure 3D**), suggesting that the engineered bacterial colonist is not simply a source of antigen and adjuvant; bacterial viability and (potentially) prolonged antigen exposure are required for the immune stimulatory response, even though no infection is mounted.

Second, antibody-mediated depletion of CD8+ T cells or all TCRβ+ cells eliminates the antitumor effect, consistent with a role for CD8+ and CD4+ T cells in the *S. epi-*OVA-induced response (**Figures 3D** and **S3**).

Finally, the subcellular localization of the antigen in *S. epidermidis* can direct a CD8+ versus CD4+ T cell response *in vivo* (**Figure 3E**). To determine the localization and antigen requirements for the antitumor effect, we colonized mice with *S. epidermidis* strains harboring different versions of OVA before injecting B16-OVA tumor cells subcutaneously into the right flank. Since *S. epi-*wOT1 only contains the CD8+ T cell antigen, we colonized mice with *S. epi-*wOVA—which contains full-length OVA—to determine whether a wall-displayed construct with CD8+ and CD4+ antigens could elicit a response. However, *S. epi-*wOVA showed no antitumor effect compared to control. In contrast, colonization with a combination of *S. epi-*wOT1 and *S. epi-*sOT2 decreased tumor size and increased IFNγ-expressing CD8+ T cells (**Figure 3F**), suggesting that the minimal requirements for antitumor efficacy are a wall-attached CD8+ antigen and a secreted CD4+ antigen. When we mismatch the localization and antigenic peptide identity by colonizing with *S. epi-*wOT2 and *S. epi-*sOT1, the antitumor effect is lost (**Figure 3E**) and IFNγ-expressing CD4+ and CD8+ T cells are not increased in the tumor-draining lymph node (**Figure 3F**). In contrast to the findings from our *in vitro* assay (**Figure 2B**), these *in vivo* data are consistent with a model in which antigen localization in the bacterial cell is critical: a wall-attached CD8+ epitope and a secreted CD4+ epitope are necessary for optimal antitumor activity. These results also suggest that antigen-specific CD4+ and CD8+ T cells are both required for the *S. epidermidis*-induced antitumor response.

### The antitumor effect of *S. epidermidis* extends to metastases outside the skin compartment

In the previous experiments, tumor cells were injected into the subcutaneous tissue of the flank. Although mice were colonized by topical application to the head, murine grooming behavior could distribute *S. epidermidis* broadly across the skin, raising the question of whether the bacterial colonist and the tumor need to be in close proximity for the induction of an antitumor immune response.

To address this question, we performed a similar experiment in the setting of metastatic melanoma using a cell line derived from B16-F10, a well-characterized (and more aggressive) variant of B16 melanoma. B16-F10-OVA cells constitutively expressing luciferase were injected intravenously rather than subcutaneously, resulting in metastases to the lungs (**Figure 4A**). Topical association with *S. epi*-OVA seven days prior to intravenous tumor cell injection slowed tumor progression substantially (**Figure 4C-E, S4**), demonstrating that the antitumor effect of *S. epi*-OVA is not restricted to skin and subcutaneous tissues. These data suggest that the antitumor effect of antigen-expressing *S. epidermidis* does not require an infection or proximity to the tumor.

**Fig. 4.**
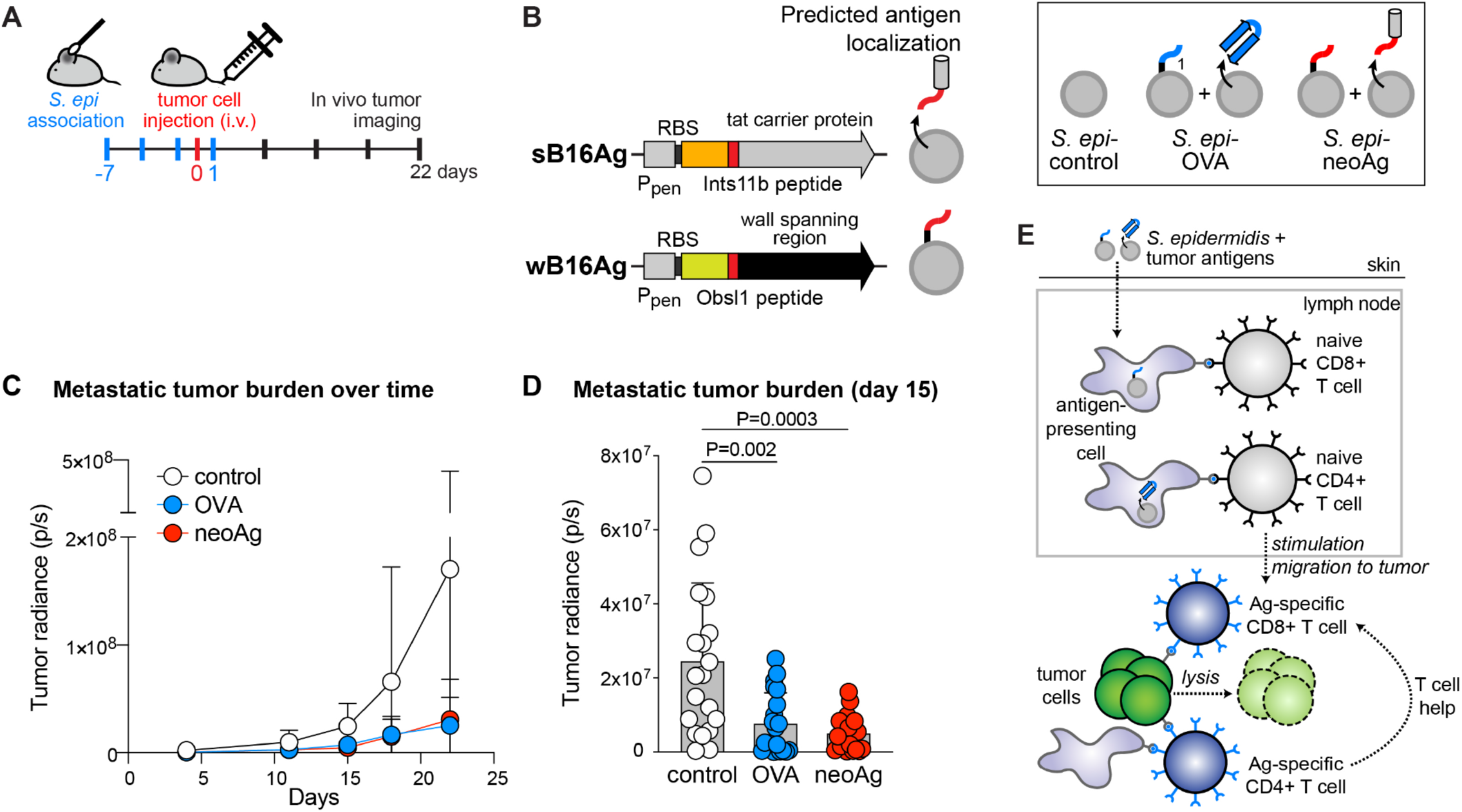
Efficacy of engineered *S. epidermidis* strains in metastatic melanoma. (A) Schematic of metastatic melanoma experiments. Mice are colonized topically with live *S. epidermidis* strains starting 7 days prior to tumor injection. On day 0, B16-F10-OVA melanoma cells (which express luciferase constitutively) are freshly prepared from growing cultures and injected intravenously into the tail vein. The tumor burden in live mice is monitored 1-2x/week by intraperitoneal luciferin injection followed by bioluminescence imaging with an IVIS Lumina Imager. Mice are sacrificed on day 22. (B) Schematic of neoantigen expression constructs and their predicted subcellular localization within *S. epidermidis*. The wall-attachment and secretion scaffolds are identical to those for wOT1 and sOT1. The neoantigen coding sequence (red box) encodes 27-aa peptides centered around Obsl1(T1764M) for the wall-attached construct (wB16Ag) or around Ints11(D314N) for the secreted construct (sB16Ag). (C) Quantification of tumor bioluminescence with dots showing the average measurement at each timepoint. (D) Bar graphs showing tumor bioluminescence on day 15 with each dot representing the measurement for each individual mouse. (E) Model of antitumor response induced by engineered commensals. Antigen-expressing strains of *S. epidermidis* colonize the skin and induce antigen-presenting cells to stimulate CD8+ or CD4+ T cells, respectively. Activated antigen-specific T cells then traffic to the tumor to restrict tumor growth.

### *S. epidermidis* producing a native neoantigen slows melanoma progression

Model antigens are useful for studying the specificity of an adaptive immune response, but their efficient processing in antigen-presenting cells and high expression in syngeneic tumor cell lines raises the question of whether this approach would work in the more realistic setting of a neoantigen naturally present in a tumor. To address this question, we engineered *S. epidermidis* to express two neoantigen-containing peptides naturally present in B16-F10 melanoma cells and previously reported to drive an antitumor response when formulated as an mRNA vaccine^44^ (**Figure 4B**). The neoantigen peptide from Obsl1(T1764M) stimulates CD8+ T cells preferentially, so we spliced a 27-aa peptide centered around the mutated residue into the wall-attachment scaffold, yielding strain *S. epi*-wB16Ag. The other neoantigen peptide, Ints11(D314N), primarily stimulates CD4+ T cells, so we spliced a 27-aa peptide harboring the mutation into a scaffold for Tat-mediated secretion, generating strain *S. epi*-sB16Ag.

We colonized mice with a mixture of *S. epi*-wB16Ag and *S. epi*-sB16Ag (termed ‘*S. epi-*neoAg’) and then injected the mice intravenously with B16-F10-OVA-luc cells seven days later. In contrast to *S. epi*-control, which failed to reduce tumor size, *S. epi*-neoAg restricted tumor growth at a comparable level to *S. epi*-OVA (**Figure 4C-E, S4**). Mice colonized by *S. epi*-neoAg do not exhibit any symptoms of autoimmunity, consistent with a model in which *S. epidermidis-*induced T cells are selective for tumor cells over healthy tissue. These data suggest that commensal-induced T cells can be redirected against a potentially broad range of host antigens.

## DISCUSSION

Our findings are consistent with a model in which an engineered commensal induces antigen-specific T cells (**Figure 4F**). Once they are primed by antigen, possibly in the skin-draining lymph nodes, these T cells can migrate to the tumor and kill tumor cells. Thus, we have co-opted the barrier response to a commensal and redirected it against a tumor, protecting the host against local and metastatic tumor progression.

Our approach—using a commensal microbe as the adjuvant and colonization as the mode of delivery in a tumor vaccine—differs from previous approaches in important ways. It does not result in an infection or require intratumoral delivery, so it is safer, simpler, and more specific than approaches that require inflammation or tissue infiltration for antitumor activity. Additionally, heat-killed *S. epi-*OVA fails to elicit a response, so we are not simply administering a purified antigen and adjuvant. The need for live bacteria suggests that *S. epidermidis* engages the immune system’s powerful (if incompletely understood) ‘barrier program’, in which the host pre-emptively develops an adaptive immune response against microbial colonists. A strain that colonizes stably may lead to prolonged antigen exposure—the equivalent of a ‘prime’ and a constant ‘boost’—and, as a result, a robust memory immune cell response.

The immune response we elicit is complex and controllable. Engineered strains of *S. epidermidis* induce a combination of antigen-specific CD8+ and CD4+ T cells; both are required for antitumor activity, consistent with recent work in the context of a neoantigen vaccine^45^. Moreover, by expressing antigens in different compartments of the bacterial cell, we can independently control the specificity of CD8+ and CD4+ T cell responses, a powerful capability that could be used to drive multifaceted responses against multiple antigens in distinct tissues.

Two improvements in design could make our approach more efficacious. First, although we observed efficacy without the need for adjuvant checkpoint blockade or cytokine therapy, combining antigen-expressing S. epidermidis strains with, e.g., antibodies targeting PD-1 or CTLA-4 could yield even more robust responses. Second, most of our experiments targeted one or two antigens in the tumor, leaving open the possibility of T cell escape by downregulating or mutating the antigen. Neoantigen vaccines typically use a small library of antigens; adapting a similar approach here would be straightforward and could improve efficacy and limit the possibility of antigen escape.

Finally, two results show the potential generality of our approach. First, engineered *S. epidermidis* protects against the growth of metastatic melanoma, so there is no need for physical proximity between the bacterium and the tumor. As a result, engineered commensal vaccines may be well suited to solid tumors and have a chance of working in a variety of tumors to which T cells have access. Second, the efficacy of neoantigen-expressing strains of *S. epidermidis* shows that our approach is not limited to model antigens; any immunogenic tumor antigen could work.

More broadly, it might be possible to engineer antigen expression into other commensal bacterial strains to elicit a wide range of antigen-specific immune cell responses. The barrier response to commensal bacteria consists of multiple adaptive immune cell types that are induced simultaneously and work together. Understanding how to redirect each one may open the door to immunotherapies for other diseases.

## Supporting information

Supplementary Materials including Methods and Supplemental Figures

## ACKNOWLEDGMENTS

We are deeply indebted to members of the Fischbach Group for helpful suggestions and comments on the manuscript. We thank Yasmine Belkaid and members of her lab for useful discussions. We thank the Stanford animal facility staff for help with animal husbandry. Cell sorting and flow cytometry analyses were performed on instruments in the Stanford Shared FACS Facility with help from M. Weglarz. *S. epidermidis* strain NIHLM087 was a gift from Julie Segre and Yasmine Belkaid, NIH. pMS182 (pLI50-Ppen-GFP-mut2) was a gift from Suzanne Walker, Harvard University. B16-F0-OVA was a gift from Nathan Reticker-Flynn from the lab of Edgar Engleman, Stanford University. This work was supported by the Stanford Microbiome Therapies Initiative, an HHMI Hanna H. Gray Fellowship (Y.E.C.); an HHMI-Simons Faculty Scholar Award (M.A.F.); a Fellowship for Science and Engineering from the David and Lucile Packard Foundation (M.A.F.); an Investigators in the Pathogenesis of Infectious Disease award from the Burroughs Wellcome Foundation (M.A.F.); NIH grant DK110174 (M.A.F.); the Chan Zuckerberg Biohub (M.A.F.); the Human Frontier Science Program LT000493/2018-L (K.N.) and the Fellowship of Astellas Foundation for Research on Metabolic Disorders (K.N.).

## AUTHOR CONTRIBUTIONS

Y.E.C., K.N., and M.A.F. conceived and designed the experiments. Y.E.C., K.A., and A.D. performed the experiments. Y.E.C. and M.A.F. analyzed data and wrote the manuscript. All authors discussed the results and commented on the manuscript.

## SUPPLEMENTARY MATERIALS

Materials and Methods

Figures S1-S3

Table S1

