## Supplementary Materials including Methods and Supplemental Figures for "Eliciting a potent antitumor immune response by expressing tumor antigens in a skin commensal"

### SUPPLEMENTARY MATERIALS for
### Eliciting a potent antitumor immune response by expressing tumor antigens in a skin commensal

Y. Erin Chen[1,2,3,5], Katayoon Atabakhsh[1,2,3], Alex Dimas[1,2,3], Kazuki Nagashima[1,2,3], Michael A. Fischbach[1,2,3,4]*

[1]Department of Bioengineering, Stanford University, Stanford, CA 94305, USA
[2]Department of Microbiology & Immunology, Stanford University School of Medicine, Stanford, CA 94305, USA
[3]ChEM-H Institute, Stanford University, Stanford, CA 94305, USA
[4]Chan Zuckerberg Biohub, Stanford, CA 94305, USA
[5]Dermatology Service, San Francisco Veterans Administration Health Care System, San Francisco, CA, USA

**This PDF file includes:**

Materials and Methods
Figs. S1 to S3
Table S1

#### MATERIALS AND METHODS

##### Genetic engineering of *Staphylococcus epidermidis*

*Staphylococcus epidermidis* LM087 was obtained from the NIH and grown in Difco Brain Heart Infusion media (BD 237200) at $37^{o}$C with shaking. Prior to liquid growth, individual colonies were selected by growth overnight on Difco Brain Heart Infusion agar plates (BD 241810). Transformation of *S. epidermidis* was performed as previously described[1]. All *S. epidermidis* strains harboring a plasmid were grown in BHI liquid media or on BHI agar plates containing chloramphenicol 10 μg/ml.

To generate electrocompetent cells, *S. epidermidis* was grown in BHI media with 0.5 M sorbitol (BHIS) overnight at $37^{o}$C with shaking and then back-diluted to an OD of 0.15 to 0.25. Back-diluted cultures were grown in BHIS until an OD of 0.7 to 0.9 at $37^{o}$C with shaking (approximately 2 hours). At this time, cultures were placed on ice and then pelleted at 3500xg for 10 minutes at $4^{o}$C. Cultures were resuspended in equal volume 10% ice-cold glycerol. Spin and resuspension steps were repeated for a total of four to five 10% glycerol washes. Each 50 ml of culture OD 0.7 to 0.9 was resuspended in 100 ul of 10% ice cold glycerol and used for one to two electroporation reactions. Competent cells can be saved at $-80^{o}$C. Upon thawing on ice, heat shock time (detailed below) is reduced to 1 minute. Freeze-thawed competent cells are not as competent as fresh competent cells but can be used if the plasmid preparation is of sufficiently high quality and concentration.

Approximately 0.5-1 μg of plasmid isolated from DC10B *E. coli* or 0.5-1 μg of minicircle plasmid from JCM2 *E. coli*[2] was added to 100 μl of competent *S. epidermidis* and cells. Method A works for strains LM018, LM061, LM087, LM088, LM095, NIH05001, ATCC12228, SK135, W23144, KPL1815. Method B works for strains ATCC35984 (RP62A), DSM20042, BCM0060. For method A, cells and plasmid in 10% glycerol were heat-shocked at $56^{o}$C for 2 minutes, then immediately transferred to a 0.1 cm cuvette, electroporated, and transferred to 3 ml of room temperature BHIS. For method B, cells and plasmid in 10% glycerol were pre-warmed at room temperature for 5 minutes, electroporated in a 0.1 cm cuvette, diluted into 1 ml of BHIS prewarmed to $56^{o}$C to heat-shock for 2 minutes, and then diluted with 3 ml of room temperature BHIS. For both methods, the electroporation program was 1.8 kV, 1 pulse, with a typical time constant of 2.3-2.5 msec using a Bio-Rad Micropulser.

Following heat shock and electroporation, cells in BHIS were recovered at $37^{o}$C for 1.5-2 hours if using a replicative plasmid and at $28^{o}$C for 4 hours if using a temperature-sensitive plasmid. After recovery, cells were spun down at 3500 x g for 10 minutes and plated onto BHIS plates with chloramphenicol (10 μg/ml).

**Design of *Staphylococcus epidermidis* antigen-expression constructs**

All antigen expression constructs were made on the plasmid backbone pLI50-Ppen-GFP-mut2 (also named pMS182, gift from Suzanne Walker, Harvard University)[3]. All primer and gBlock sequences and plasmid names are listed in Table S1. All primers and gBlocks were ordered from IDT and all coding sequences were optimized for *Staphylococcus* using the IDT codon optimization tool.

The plasmid backbone and promoter were linearized by PCR with primers 199 and 223, yielding linearized pLI50-Ppen. To create pLI50-Ppen-cOVA, gBlock 474 (containing the ribosome binding site from *S. aureus* gene *hld* followed by the coding sequence for OVA) was inserted into linearized pLI50-Ppen by Gibson assembly (NEB E2611L).

To create pLI50-Ppen-sOVA, the following pieces were assembled together by Gibson assembly: (1) gBlock 479 containing the N-terminal Tat secretion signal from *S. aureus fepB* (gene locus name SAOUHSC_00326), (2) gBlock 474 (containing OVA) amplified with primers 475 and 476 to remove the N-terminal methionine, and (3) linearized pLI50-Ppen containing the plasmid backbone.

To create pLI50-Ppen-sOT1, pLI50-Ppen-sOVA3pep was initially created by assembling the following pieces together: (1) gBlock 508 containing OVA3pep and the secretion signal, (2) the *fepB* carrier protein (SAOUHSC_00326) amplified by primers 511 and 512 from *S. aureus* strain NCTC 8325, and (3) linearized pLI50-Ppen. Then, to create pLI50-Ppen-sOT1, primers 693 and 694 were used to amplify pLI50-Ppen-sOVA3pep and simultaneously replace OVA3pep with the OT1 peptide.

To create pLI50-Ppen-sOT2, primers 701 and 702 were used to amplify pLI50-Ppen-sOT1 and simultaneously replace the OT1 peptide with the OT2 peptide.

To create pLI50-Ppen-wOVA, primers 480 and 481 were used to amplify the *S. aureus* protein A sortase signal sequence from gBlock 341, primers 482 and 483 were used to amplify OVA from gBlock 474 and remove the N-terminal methionine, and primers 371 and 484 were used to amplify the wall-spanning protein A XM domain from gBlock 342[4]; these pieces were assembled into linearized pLI50-Ppen. pLI50-Ppen-wOT1 was constructed similarly, except primers 480 and 485 were used to PCR gBlock 341 (N-terminal sortase signal sequence) and primers 486 and 484 were used to PCR gBlock 342 (XM domain); these primers simultaneously amplified the wall-attachment scaffold and added the OT1 peptide for assembly into linearized pLI50-Ppen.

To construct pLI50-Ppen-wOT2, overlapping primers 699 and 700 were used to amplify the entire pLI50-Ppen-wOT1 plasmid and simultaneously replace the OT1 peptide with the OT2 peptide.

To construct pLI50-Ppen-wOVA3pep, OVA3pep was amplified from pLI50-Ppen-sOVA3pep with primers 572 and 573 and then assembled into pLI50-Ppen-wOT1 linearized with primers 371 and 481.

To generate the wall-attached B16 neoantigen construct (wB16Ag), gBlock 723 containing the 27-aa neoantigen peptide from Obsl1(T1764M) was assembled into pLI50-Ppen-wOT1 linearized with primers 371 and 634. To generate the secreted B16 neoantigen construct (sB16Ag), gBlock 716 containing the 27-aa neoantigen peptide from Ints11(D314N) was assembled into pLI50-Ppen-sOVA linearized with primers 717 and 223. Immunogenicity of these neoantigens have been previously described[5,6].

**Western blot**

OVA-expressing *S. epidermidis* strains were grown in BHI overnight and normalized by $OD_{600}$. The equivalent of 1 ml of $OD_{600}$ = 10 bacteria was pelleted, resuspended in 1 ml PBS, and then lysed by bead beating using 0.1 mm beads and the Tissue Lyser II (Qiagen 85300) at room temperature for 25 minutes. The resultant lysed sample was briefly spun down, added to sample buffer with DTT (NEB B7703S) and boiled 10 min at 95$^{o}$C before loading onto a protein gel. For immunoblotting of secreted protein, 1 ml of $OD_{600}$ = 10 supernatant from overnight cultures was incubated with 250 ul of 50% trichloroacetic acid (w/v) for 20 minutes on ice. Precipitated proteins were pelleted at 18000g x 10 minutes at 4$^{o}$C. Pellets were washed with 250 ul ice cold acetone twice, dried for 2-5 minutes at 95$^{o}$C, and then solubilized in 100 $\mu$l protein sample buffer. Samples were then loaded onto a NuPAGE 4-12% Bis-Tris gel (Invitrogen NP0321PK2) with MES running buffer, and protein separation was achieved by running at 105 V for 2 hours. Transfer was done at 150V in 4$^{o}$C for 1 hour using 0.2 $\mu$m PVDF and FLASHBlot transfer buffer (VWR 103254-862). Immunoblotting was performed using polyclonal rabbit anti-OVA antibody 1:500 (Creative Diagnostics DPAB27549), secondary goat anti-rabbit HRP antibody 1:2000 (Invitrogen 656120), and imaged following exposure to SuperSignal West Pico PLUS Chemiluminescent Substrate (Thermo Scientific 34580) per manufacturer's instructions.

**Cell lines and tissue culture**

All mouse B16 melanoma lines were grown in Dulbecco's modified Eagle's medium (DMEM, ATCC 30-2002) supplemented with 10% (v/v) fetal bovine serum. Cell growth was maintained at 37°C in a 5% $CO_2$ atmosphere, passaging at a 1:10 dilution every 2-3 days. B16-F0-OVA was provided by Nathan Reticker-Flynn from the laboratory of Edgar Engleman, Stanford University. B16-F10 producing luciferase was obtained from ATCC (B16-F10-Luc2, CRL-6475-LUC2). B16-F10-OVA was made by lentiviral transduction into B16-F10-Luc2. Lentivirus was produced by transfecting HEK 293T cells with packaging plasmids pCMV-ΔR8.91[7] and pMD2.G (pMD2.G was a gift from Didier Trono (Addgene plasmid # 12259)) and a cargo plasmid expressing OVA-T2A-GFP behind the EF-1α promoter, amplified from the plasmid pU6-sgRNA Ef1alpha Puro-T2A-GFP (gift from Jonathan Weissman, Addgene plasmid #111596)[8]. Production of lentivirus carrying the OVA-GFP cargo plasmid was performed as previously described using HEK 293T cells[9]. To select for B16-F10-OVA clones that stably and highly express OVA, cells were sorted twice for high GFP expression 4 days and 14 days after viral transduction. High-expressing clones were expanded and frozen for future use. After thawing and expansion of frozen aliquots and immediately before injection into mice, B16-F10-OVA cells were confirmed to have high OVA-GFP expression using flow cytometry.

**In vivo tumor models**

Six-to-ten-week-old female C57BL/6 mice were purchased from Taconic Biosciences. Subcutaneous and metastatic tumor experiments were performed as previously described[10]. Briefly, for subcutaneous tumor experiments, B16-F0-OVA cells were grown as detailed above, freshly harvested when approximately 50% confluent, and resuspended in PBS. $1.25 \times 10^5$ cells in 200 μl per mouse were injected subcutaneously into the right flank on day 0. For metastatic tumor experiments, $2 \times 10^5$ B16-F10-OVA cells suspended in 200 μl PBS were delivered intravenously on day 0. Bioluminescence of tumors was measured 1-2 times per week. Bioluminescence was measured by injecting 3 mg per 20 g mouse of D-luciferin (Biosynth International, L-8220) intraperitoneally and imaged at 90 to 180 seconds range of exposure using the IVIS Lumina Imager (PerkinElmer). Data was analyzed using the Living Image Software (PerkinElmer) and tumor burden was reported as total radiance over the entire mouse body.

Topical associations with *S. epidermidis* strains were performed as previously described on days -7, -4 or -5, -2, and +1 using cultures normalized to $OD_{600}$ = 3 with BHI media[11].

For T cell depletion experiments, mice were treated with intraperitoneal injections of 200 μg/mouse anti-CD8α (2.43, BioXCell BE0061) or 200 μg/mouse anti-TCRβ (H57-597, BioXCell

BE0102) neutralizing antibodies as previously described[12,13]. Injections were started 4 days prior to tumor injection and continued twice weekly until 4 days prior to sacrifice.

All animal procedures were performed in accordance with guidelines established by Stanford university institutional animal care and use committee guidelines.

**Mixed lymphocyte assay**

Eight-to-twelve-week-old female CD45.1, OT-I, and OT-II mice were purchased from Jackson Laboratories. Spleens were dissociated as detailed above.

Dendritic cells were isolated from spleens of CD45.1 mice that were pre-treated with an intraperitoneal injection of B16 producing Flt3L to increased dendritic cell yield[14]. Mice were sacrificed approximately 10-14 days after injection and dendritic cells were isolated from the spleens using the CD11c microbeads (Miltenyi 130-125-835) and manufacturer's instructions. $1 \times 10^6$ CD45.1 dendritic cells were co-cultured in DMEM with 10% FBS with one of the following stimulants for 2 hours at 37°C in a 5% $CO_2$ atmosphere: 10 μl of PBS, eBioscience Cell Stimulation Cocktail (Thermo Fisher 00-4975-93), 10 μl of ovalbumin (10 mg/ml in PBS), 10 μl of OT-1 or OT-2 peptide (Anaspec, 1 mg/ml in PBS), or 10 μl of heat-shocked bacteria resuspended in PBS. *S. epidermidis* strains were pelleted and resuspended in PBS to $OD_{600}$ = 1 and then 1 ml was heat-shocked in a bead bath at 70°C for 30 minutes and frozen into single use aliquots at -20°C, then thawed on ice prior to use.

After 2 hours, $2.5 \times 10^4$ OT-I or OT-II T cells were added and co-cultured for another 4 hours before staining and fixation as detailed below. T cells were isolated from spleens of OT-I and OT-II mice using the Pan T Cell Isolation Kit II (Miltenyi 130-095-130) and manufacturer's instructions.

**Flow cytometry**

Spleens were dissociated using the GentleMACS Spleen Dissociation Kit (Miltenyi 130-095-926) and manufacturer's instructions to generate single cell suspensions. Tumor-draining inguinal lymph nodes were digested for 20 minutes at 37°C in RPMI supplemented with 0.25 mg/ml Liberase TL enzyme blend (Sigma 5401020001) and then mashed through a 100 μm cell strainer to generate single cell suspensions, as previously described[15].

For detection of basal cytokine potential, single cell suspensions were cultured directly ex vivo in a 96-well U-bottom plate in RPMI 1640 supplemented with 10% fetal bovine serum (FBS) and stimulated with 50 ng/ml phorbol myristate acetate (PMA) (Sigma-Aldrich) and 5 μg/ml ionomycin (Sigma-Aldrich) in the presence of brefeldin A (GlogiPlug, BD Biosciences) for 2.5 hours at 37°C in 5% CO2 as previously described[15].

Single cell suspensions were incubated with combinations of the following fluorophore-conjugated antibodies against surface markers: CD4 (RM4-5), CD8β (eBioH35-17.2), CD45.1 (A20), CD45.2 (104), CD90.2 (53-2.1), TCRβ (H57-597) in Hank's buffered salt solution (HBSS) for 20 min at 4°C and then washed. Fixable Viability Dye eFluor 780 (eBioscience 65-0865-18) was used to exclude dead cells. Cells were then fixed for 60 min at 4°C or 30 min at room temperature using the fixation/permeabilization buffer supplied with the Transcription Factor Staining set (eBioscience 00-5523-00) and then washed. For intracellular staining, cells were then stained with the combinations of the following fluorochrome-conjugated antibodies in permeabilization buffer for 1 hour at 4°C: IFN-γ (XMG-1.2), IL-17A (eBio17B7), Foxp3 (FJK-16s), and Nur77 (12.14). All staining was performed in the presence of purified anti-mouse CD16/32 (clone 93) and 0.2 mg/ml purified rat gamma globulin (Jackson Immunoresearch). All antibodies were purchased from eBioscience (Life Technologies), Biolegend, or BD Biosciences. Cell acquisition was performed on an LSR II flow cytometer using FACSDiVa software (BD Biosciences) and data analyzed using FlowJo software (TreeStar).

**Statistical Analyses**

All statistical analysis was done with the software Graphpad Prism (GraphPad Software, La Jolla, CA). Student's t-test was used whenever appropriate. P values less than 0.05 were considered statistically significant.

Figure S1

**Western blot of OVA-expressing *S. epi***

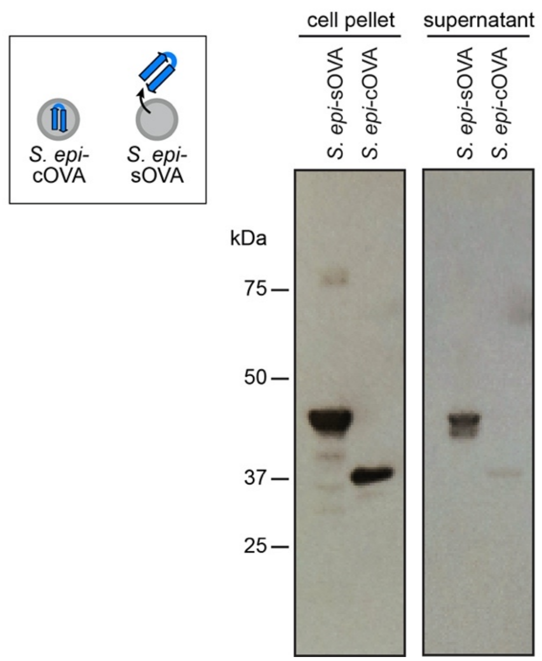

**Fig. S1. Western blot of engineered *S. epidermidis* strains.** Western blot of protein extracted from cell pellets or overnight liquid culture supernatants of *S. epi*-sOVA or *S. epi*-cOVA.

Figure S2

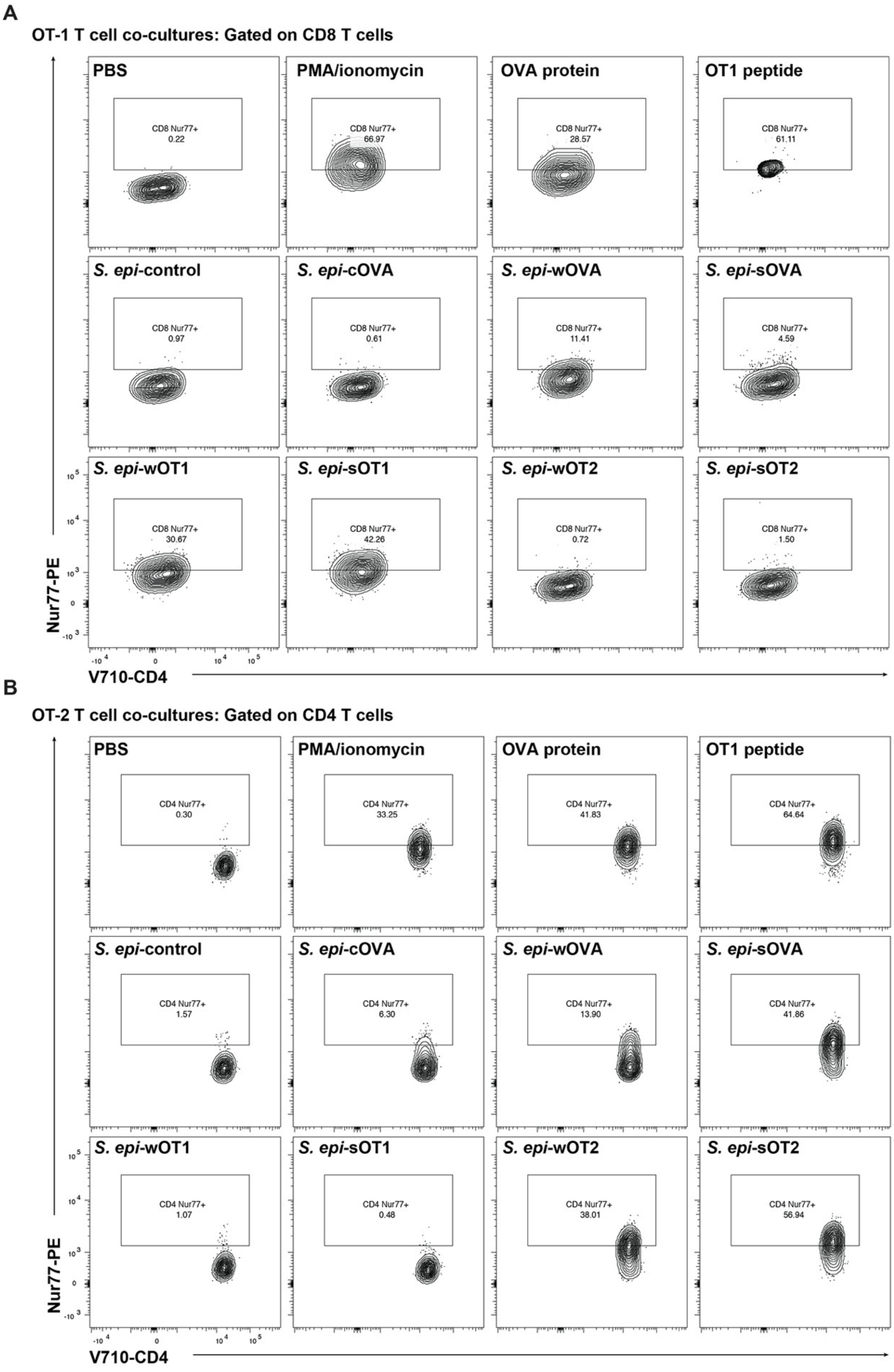
A
OT-1 T cell co-cultures: Gated on CD8 T cells
PBS
CD8 Nur77+
0.22
PMA/ionomycin
CD8 Nur77+
66.97
OVA protein
CD8 Nur77+
28.57
OT1 peptide
CD8 Nur77+
61.11
S. epi-control
CD8 Nur77+
0.97
S. epi-cOVA
CD8 Nur77+
0.61
S. epi-wOVA
CD8 Nur77+
11.41
S. epi-sOVA
CD8 Nur77+
4.59
S. epi-wOT1
CD8 Nur77+
30.67
S. epi-sOT1
CD8 Nur77+
42.26
S. epi-wOT2
CD8 Nur77+
0.72
S. epi-sOT2
CD8 Nur77+
1.50
Nur77-PE
V710-CD4
B
OT-2 T cell co-cultures: Gated on CD4 T cells
PBS
CD4 Nur77+
0.30
PMA/ionomycin
CD4 Nur77+
33.25
OVA protein
CD4 Nur77+
41.83
OT1 peptide
CD4 Nur77+
64.64
S. epi-control
CD4 Nur77+
1.57
S. epi-cOVA
CD4 Nur77+
6.30
S. epi-wOVA
CD4 Nur77+
13.90
S. epi-sOVA
CD4 Nur77+
41.86
S. epi-wOT1
CD4 Nur77+
1.07
S. epi-sOT1
CD4 Nur77+
0.48
S. epi-wOT2
CD4 Nur77+
38.01
S. epi-sOT2
CD4 Nur77+
56.94
Nur77-PE
V710-CD4

**Fig. S2. Antigen-specific T cell stimulation by engineered strains of *S. epidermidis*.** Representative flow plots of data from the mixed lymphocyte assay shown in Figure 2B. (A) Live CD45.1- CD90.2+ TCRβ+ CD8β+ OT-I T cells are analyzed for Nur77 expression after co-culture with dendritic cells and control stimulants (top row) or engineered *S. epidermidis* strains. (B) Live CD45.1- CD90.2+ TCRβ+ CD4+ OT-2 T cells are analyzed for Nur77 expression after co-culture with dendritic cells and control stimulants (top row) or engineered *S. epidermidis* strains.

Figure S3

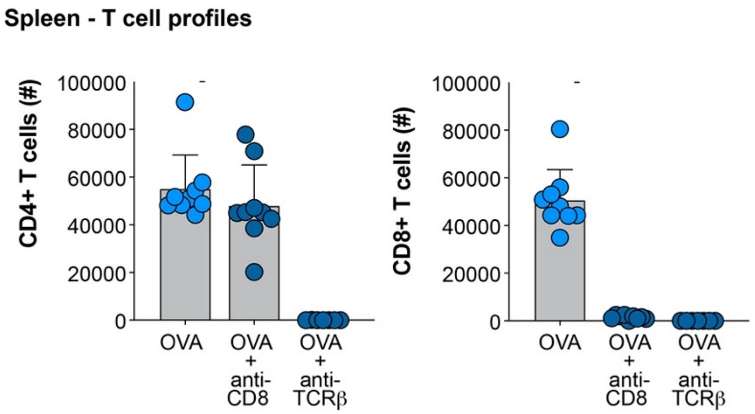

**Fig. S3. Successful T cell depletion using neutralizing antibodies.** These data serve as a control for the data shown in Figure 2D. All mice were topically associated with *S. epi*-OVA and injected with subcutaneous B16-F0-OVA tumors. In the group treated with anti-CD8 neutralizing antibody, spleens have a normal number of CD4+ T cells (left) but no CD8+ T cells (right). In the group treated with anti-TCRβ neutralizing antibody, both CD4+ (left) and CD8+ (right) T cells are depleted.

**Figure S4**

**Bioluminescence imaging of metastatic B16-F10-OVA**

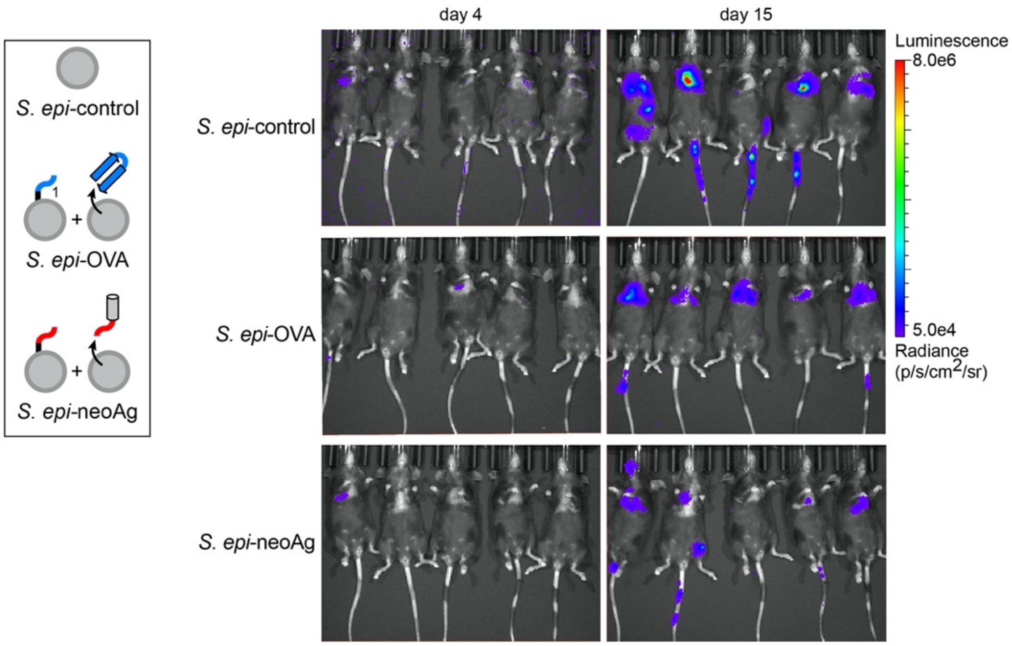

**Fig. S4. Bioluminescence imaging of metastatic B16-F10-OVA tumor burden.** Representative images showing the bioluminescence of metastatic tumors using in vivo imaging. All mice were topically associated with *S. epi*-control, *S. epi*-OVA, or *S. epi*-neoAg. These images were obtained on day 4 or day 15 after tail vein tumor injection of B16-F10-OVA (with constitutive luciferase expression).

#### Table S1. Components used for engineering *S. epidermidis*

##### Primers

| Primer number | Sequence |
|---|---|
| 199 | ATGTATATCTCCTTCTTAAATTAATTAGTT |
| 223 | CCTATTCTAAATGCATAATAAATACTG |
| 371 | AAACCTGGTAAAGAAGATGGTAACGGAG |
| 475 | GGTTCAATTGGAGCTGCGTC |
| 476 | CAGTATTTATTATGCATTTAGAATAGG |
| 480 | ATTAATTTAAGAAGGAGATATACATGGAAGGA |
| 481 | AGCATTTGCAGCAGGTGTTA |
| 482 | TAACACCTGCTGCAAATGCTGGTTCAATTGGAGCTGCGTC |
| 483 | ACCATCTTCTTTACCAGGTTTAGGACTAACACAACGTCCG |
| 484 | CAGTATTTATTATGCATTTAGAATAGGTTATAGTTCGCGACGACGT |
| 485 | CAACTTCTCAAAATTAATTATAGAAGCATTTGCAGCAGGTGTTA |
| 486 | TCTATAATTAATTTTGAGAAGTTGAAACCTGGTAAAGAAGATGGTAAC |
| 511 | ATTAACTGAGTGGAGTGGTGTTGGTAGCATGT |
| 512 | CAGTATTTATTATGCATTTAGAATAGGTTAGTCAAATAATGTTTCACCAAGGTA |
| 572 | GTAACACCTGCTGCAAATGCTCAGACTGCTATGGTGTTGGTG |
| 573 | TACCATCTTCTTTACCAGGTTTCCACTCAGTTAATTTTTCAAAGTT |
| 634 | AGCATTTGCAGCAGGTGTTACA |
| 693 | TCTATAATTAATTTTGAGAAGTTGAGTGGTGTTGGTAGCATGTGGTCTTTC |
| 694 | CAACTTCTCAAAATTAATTATAGATGCGCCAATTGCAACACCGGCACCG |
| 699 | CATGCGGAGATTAATGAGGCTGGCAAACCTGGTAAAGAAGATGGTAACGG |
| 700 | GCCAGCCTCATTAATCTCCGCATGCGCCGCATGAGCATTTGCAGCAGGTGTTAC |
| 701 | GCGCATGCGGAGATTAATGAGGCTGGCAGTGGTGTTGGTAGCATGTGGTCTTTCAAATCA |
| 702 | CCTCATTAATCTCCGCATGCGCCGCATGTGCGCCAATTGCAACACCGGCACCGCCAATAC |
| 717 | TGCGCCAATTGCAACACCGG |

##### gBlocks

| gBlock number | Sequence |
|---|---|
| 341 | ATTAATTTAAGAAGGAGATATACATGGAAGGAGTGATTTCATTGAAAAAGAAAAACATTTATTCAATTCGTAAACTAGGTGTAGGTATTGCATCTGTAA<br>CTTTAGGTACATTACTTATATCTGGTGGCGTAACACCTGCTGCAAATGCTGATTATAAGGATCACGACGGTGACTACAAGGATCACGATATTGACTAT<br>AAGGATGATGACGACAAA |
| 342 | AAACCTGGTAAAGAAGATGGTAACGGAGTACATGTCGTTAAACCTGGTGATACAGTAAATGACATTGCAAAAGCAAACGGCACTACTGCTGACAAAA<br>TTGCTGCAGATAACAAATTAGCTGATAAAAACATGATCAAACCTGGTCAAGAACTTGTTGTTGATAAGAAGCAACCAGCAAACCATGCAGATGCTAAC<br>AAAGCTCAAGCATTACCAGAAACTGGCGAAGAAAATCCATTCATCGGTACAACTGTATTTGGTGGATTATCATTAGCCTTAGGTGCAGCGTTATTAGC<br>TGGACGTCGTCGCGAACTATAA |
| 474 | AACTAATTAATTTAAGAAGGAGATATACATGGAAGGAGTGATTTCAATGGGTTCAATTGGAGCTGCGTCAATGGAATTTTGTTTCGACGTTTTTAAGG<br>AATTAAAAGTACACCATGCTAACGAGAACATATTCTATTGCCCGATTGCTATCATGTCTGCATTAGCTATGGTATATTTGGGAGCAAAGGACAGTACA<br>CGTACGCAAATAAATAAGGTTGTCCGTTTCGACAAATTGCCAGGTTTTGGAGATTCAATAGAGGCGCAGTGCGGAACAAGTGTTAATGTACATAGTT<br>CTTTACGTGATATATTGAATCAGATAACGAAACCTAACGACGTCTATTCTTTTAGTTTAGCTTCAAGATTGTACGCGGAGGAGCGTTATCCTATCTTG<br>CCAGAATATTTACAATGTGTAAAAGAATTATACCGAGGCGGATTAGAACCTATTAATTTTCAGACGGCGGCAGATCAAGCTCGTGAATTGATTAACTC<br>ATGGGTGGAAAGTCAAACTAATGGTATTATACGAAACGTCTTACAACCTAGTTCTGTCGATTCTCAGACTGCTATGGTGTTGGTGAATGCGATTGTCT<br>TTAAAGGATTATGGGAGAAAGCTTTCAAGGATGAAGACACTCAAGCAATGCCGTTCCGAGTAACAGAACAGGAATCTAAGCCGGTGCAGATGATGT<br>ATCAAATTGGCTTATTCAGAGTCGCGTCAATGGCGAGTGAAAAGATGAAAATCTTAGAGTTACCTTTTGCATCAGGTACAATGTCAATGTTGGTCTTA<br>TTGCCAGACGAAGTTTCTGGCTTAGAGCAGTTAGAATCTATAATTAATTTTGAGAAGTTGACAGAGTGGACGTCAAGTAATGTCATGGAGGAACGTA<br>AGATCAAGGTCTACTTACCACGTATGAAAATGGAGGAAAAGTATAACTTAACGTCAGTCTTAATGGCGATGGGCATCACGGACGTCTTTTCTTCTAGT<br>GCGAACTTATCTGGTATATCTAGTGCGGAGTCTTTAAAAATATCTCAAGCAGTCCATGCGGCGCATGCGGAGATTAATGAGGCTGGCCGTGAAGTA<br>GTAGGTTCTGCGGAAGCGGGAGTAGATGCAGCTTCAGTCTCAGAGGAGTTTAGAGCTGACCACCCATTCTTGTTCTGTATCAAACACATAGCAACTA<br>ACGCTGTCTTGTTCTTCGGACGTTGTGTTAGTCCTTGACCTATTCTAAATGCATAATAAATACTG |
| 479 | AACTAATTAATTTAAGAAGGAGATATACATGGAAGGAGTGATTTCAATGACAAATTATGAACAAGTTAACGATAGTACGCAATTTTCAAGACGTACATT<br>TTTGAAAATGTTAGGTATTGGCGGTGCCGGTGTTGCAATTGGCGCAGGTTCAATTGGAGCTGCGTC |
| 508 | AACTAATTAATTTAAGAAGGAGATATACATGGAAGGAGTGATTTCAATGACAAATTATGAACAAGTTAACGATAGTACGCAATTTTCAAGACGTACATT<br>TTTGAAAATGTTAGGTATTGGCGGTGCCGGTGTTGCAATTGGCGCACAGACTGCTATGGTGTTGGTGAATGCGATTGTCTTTAAAGGATTATGGCCT<br>TTTGCATCAGGTACAATGTCAATGTTGGTCTTATTGCCAGACGAAGTTTCTGGCTTAGAACAGTTAGAATCAATCATAAACTTTGAAAAATTAACTGAG<br>TGGAGTGGTGTTGGTAGCATGT |

| | |
|---|---|
| 716 | TGGCGGTGCCGGTGTTGCAATTGGCGCACCTGAGATAAGAGTTACGCCATTAGGCGCGGGCCAAGACGTGGGCAGAAGTTGCATATTAGTTAGTA<br>TAAGTGGTAAAAACGTGATGTTAGATTGCGGTATGCACATGGGTTACAATGACGATAGACGTTTCCCGGATTTTTCATATATTACACAGTCTGGCAGA<br>TTAACAGACTTTTTGGACTGTGTTATAATAAGTCATTTCCACTTAGACCACTGTGGTGCATTGCCATACTTCTCTGAAATGGTGGGTTATGACGGTCC<br>TATATACATGACACATCCGACGCAGGCTATCTGTCCTATTTTATTAGAGGACTATAGAAAGATAGCGGTCGATAAAAAAGGCGAAGCGAATTTTTTCA<br>CTAGTCAAATGATTAAAGACTGTATGAAGAAAGTAGTCGCAGTGCACTTACATCAGACGGTTCAAGTTGATGACGAGTTGGAGATTAAGGCGTATTA<br>CGCAGGACACGTTTTGGGTGCAGCTATGTTCCAGATAAAGGTAGGTTCTGAGTCTGTCGTGTATACGGGTGATTACAACATGACTCCGGATAGACA<br>CTTGGGAGCGGCATGGATCGATAAGTGTAGACCAAACTTATTGATCACGGAGTCTACTTACGCAACGACAATTAGAGATTCAAAACGTTGCAGAGAG<br>CGAGATTTCTTGAAGAAAGTACACGAAACAGTTGAAAGAGGAGGTAAGGTCTTGATTCCAGTCTTCGCTTTAGGAAGAGCTCAAGAGTTATGCATCT<br>TGTTAGAGACGTTTTGGGAAAGAATGAATTTAAAAGTGCCAATATACTTTTCAACTGGCTTGACTGAGAAAGCGAACCATTATTACAAGTTATTTATTA<br>CATGGACAAATCAAAAGATCAGAAAGACTTTTGTACAGCGTAACATGTTCGAGTTTAAGCATATAAAAGCTTTTGATAGAACGTTTGCGAACAATCCA<br>GGACCTATGGTAGTCTTTGCAACTCCGGGTATGTTACACGCAGGCCAGTCATTGCAAATCTTCCGTAAGTGGGCGGGCAACGAAAAAAATATGGTC<br>ATTATGCCGGGATACTGCGTGCAGGGTACAGTAGGCCATAAGATATTATCAGGTCAACGAAAGTTAGAAATGGAGGGTAGACAAATGTTGGAAGTG<br>AAAATGCAAGTAGAATACATGAGTTTTTCAGCTCATGCTGATGCTAAGGGTATAATGCAGTTAGTGGGCCAGGCGGAACCAGAGTCAGTATTATTAG<br>TCCACGGAGAAGCGAAAAAGATGGAGTTTTTGCGTCAGAAAATTGAACAGGAGTTTCGAGTATCTTGTTACATGCCAGCTAATGGCGAAACTGTCAC<br>GTTACCTACGAGTCCATCAATCCCGGTTGGTATCTCATTAGGTTTGTTGAAGCGAGAGATGGTGCAGGGATTATTACCAGAGGCTAAGAAGCCTCG<br>ATTGTTACATGGCACATTGATAATGAAAGACAGTAACTTCAGATTGGTCTCTTCAGAGCAAGCGTTGAAAGAATTGGGATTAGCAGAGCATCAATTAA<br>GATTCACATGCCGAGTACACTTACAGGATACGCGAAAGGAACAGGAGACTGCTTTGCGTGTTTATTCTCATTTAAAGTCTACTTTAAAAGACCATTGC<br>GTTCAACACTTACCAGACGGCAGTGTGACTGTCGAGAGTATCTTAATTCAGGCTGCAGCTCATTCAGAAGACCCGGGCACAAAAGTGTTGTTAGTAT<br>CTTGGACTTATCAGGATGAAGAGTTAGGAAGTTTCTTAACGACGTTGTTGAAAAACGGATTGCCGCAGGCGCCGTCTTGACCTATTCTAAATGCATA<br>ATAAATACTGATAACATCTTATATTTTG |
| 723 | TTGGGTACATTGTTAATTAGTGGTGGTGTAACACCTGCTGCAAATGCTCGAGAAGGAGTCGAGTTGTGTCCAGGCAACAAATATGAGATGCGTCGA<br>CATGGTACGACACACTCTTTAGTCATACACGACAAACCTGGTAAAGAAGATGGTAACGGAGTACATGTCGTTAAACCTGGTG |

#### Plasmids and strains

| **Strain name** | **Plasmid name** |
|---|---|
| *S. epi*-cOVA | pLI50-Ppen-cOVA |
| *S. epi*-sOVA | pLI50-Ppen-sOVA |
| *S. epi*-sOT1 | pLI50-Ppen-sOT1 |
| *S. epi*-sOT2 | pLI50-Ppen-sOT2 |
| *S. epi*-sOVA3pep | pLI50-Ppen-sOVA3pep |
| *S. epi*-wOVA | pLI50-Ppen-wOVA |
| *S. epi*-wOT1 | pLI50-Ppen-wOT1 |
| *S. epi*-wOT2 | pLI50-Ppen-wOT2 |
| *S. epi*-wOVA3pep | pLI50-Ppen-wOVA3pep |
| *S. epi*-wB16Ag | pLI50-Ppen-wB16Ag |
| *S. epi*-sB16Ag | pLI50-Ppen-sB16Ag |
